# Convergent evidence for the molecular basis of musical traits

**DOI:** 10.1101/061358

**Authors:** Jaana Oikkonen, Päivi Onkamo, Veronika Kravtsov, Irma Järvelä, Chakravarthi Kanduri

## Abstract

To obtain aggregate evidence for the molecular basis of musical abilities and the effects of music, we integrated gene-level data from 101 published studies across multiple species including humans, songbirds and several other animals and used a convergent evidence method to prioritize the top candidate genes. Several of the identified top candidate genes like *EGR1, FOS, ARC, BDNF* and *DUSP1* are known to be activity-dependent immediate early genes that respond to sensory and motor stimuli in the brain. Several other top candidate genes like *MAPK10, SNCA, ARHGAP24, TET2, UBE2D3, FAM13A* and *NUDT9* are located on chromosome 4q21-q24, on the candidate genomic region for music abilities in humans. Functional annotation analyses showed the enrichment of genes involved in functions like cognition, learning, memory, neuronal excitation and apoptosis, long-term potentiation and CDK5 signaling pathway. Interestingly, all these biological functions are known to be essential processes underlying learning and memory that are also fundamental for musical abilities including recognition and production of sound. In summary, our study prioritized top candidate genes related to musical traits that are possibly conserved through evolution, as suggested by shared molecular background with other species.

## Introduction

Music perception and performance represent complex cognitive functions of the human brain. Research on families, twins and newborns, neurophysiological studies and more recently genomics studies have suggested that the abilities to perceive and practice music have a biological background^1–5^. Over the past decade, several studies ranging from candidate gene level to genome-wide scans have investigated the molecular basis of musical traits in humans. For example, conventional genetic approaches such as genome-wide linkage scans, association studies^3,6–8^ and CNV studies^9,10^ have shown that musical aptitude is linked to regions that contain genes affecting development and function of auditory pathway and neurocognitive processes. Studies of genome-wide RNA expression profiles have revealed that listening and performing music enhanced the activity of genes related to dopamine secretion and transport, neuronal plasticity, learning and memory^11,12^. Recently, positive selection regions associated with musical aptitude have been shown to contain genes that affect hearing, language development, birdsong and reward mechanism^13^. Despite these independent findings, aggregate evidence for the molecular basis of musical traits remains lacking.

Musical abilities in humans are based on sound perception and production that are well preserved in evolution^14–16^. Even birdsong is known to have musical features: it is a combination of rhythms, pitches and transitions that induce emotional responses^17^ and vocal learning show similar features between songbirds and humans^18^. In zebra finches, vocal learning happens during sensitivity period from 25 to 65 dph (days post-hatch)^19^. Similar sensitivity period to learn music and language has been reported in humans^20^. Convergent evolution of hearing genes has been found in echo-locating bats and dolphins^21^. Interestingly, mammals, including modern humans, have been shown to share similar inner ear structures even with insects^22^. A shared background of sound perception and production between evolutionarily as distant species as humans and songbirds has been found in our previous studies, where several homologous genes known to affect song learning and singing in songbirds were up-regulated after music perception and performance in humans^11,12^. All these evidence suggest a high evolutionary conservation, or convergent evolution, of molecular mechanisms related to sound perception and production. Therefore, data from relevant animal models like songbirds might be used as an additional layer of evidence when outlining the contours of the genetic landscape underlying musical traits in humans.

To obtain convergent evidence for the molecular basis of musical traits, we integrated gene-level data from a wide range of studies including genome-wide linkage and association studies, gene expression studies, candidate gene studies and other molecular studies in both humans and relevant animal models. A similar strategy has earlier been used successfully to identify top candidate genes underlying several neuropsychiatric diseases^23–25^. Here, we curated a database of 101 published studies with 7883 genes by integrating our own and other human studies as well as animal studies related to music and musical abilities including studies on songbirds, mice and several other species. We used the convergent evidence method (CE) implemented by one of the authors (C.K.) in GenRank Bioconductor package to prioritize the candidate genes. Functional enrichment analysis of the prioritized genes suggests that the top candidate genes are known to affect cognition, learning, memory, excitation of neurons, quantity of catecholamine and long-term potentiation.

## Results

### Study database

We retrieved a total of 326 articles related to music, of which 101 articles were shortlisted after filtering out irrelevant articles. The retained articles included biomarker and candidate gene studies and genome-wide studies at both DNA-and RNA-level across several species including humans, songbirds and mice (**Figure 1**). All the chosen studies are listed in the supplementary information and in **Tables S1-S3** and the summary statistics of the study database are shown in **Table 1**. The genes and molecules identified in animal model studies were translated into human genes through homologs. A total of 7883 genes and biomarkers were identified at least once from the 101 published studies related to musical abilities (**Table S4**). Nearly a quarter of the genes (1755 genes) were found within linkage regions (**Figure 2**).

**Figure 1.**
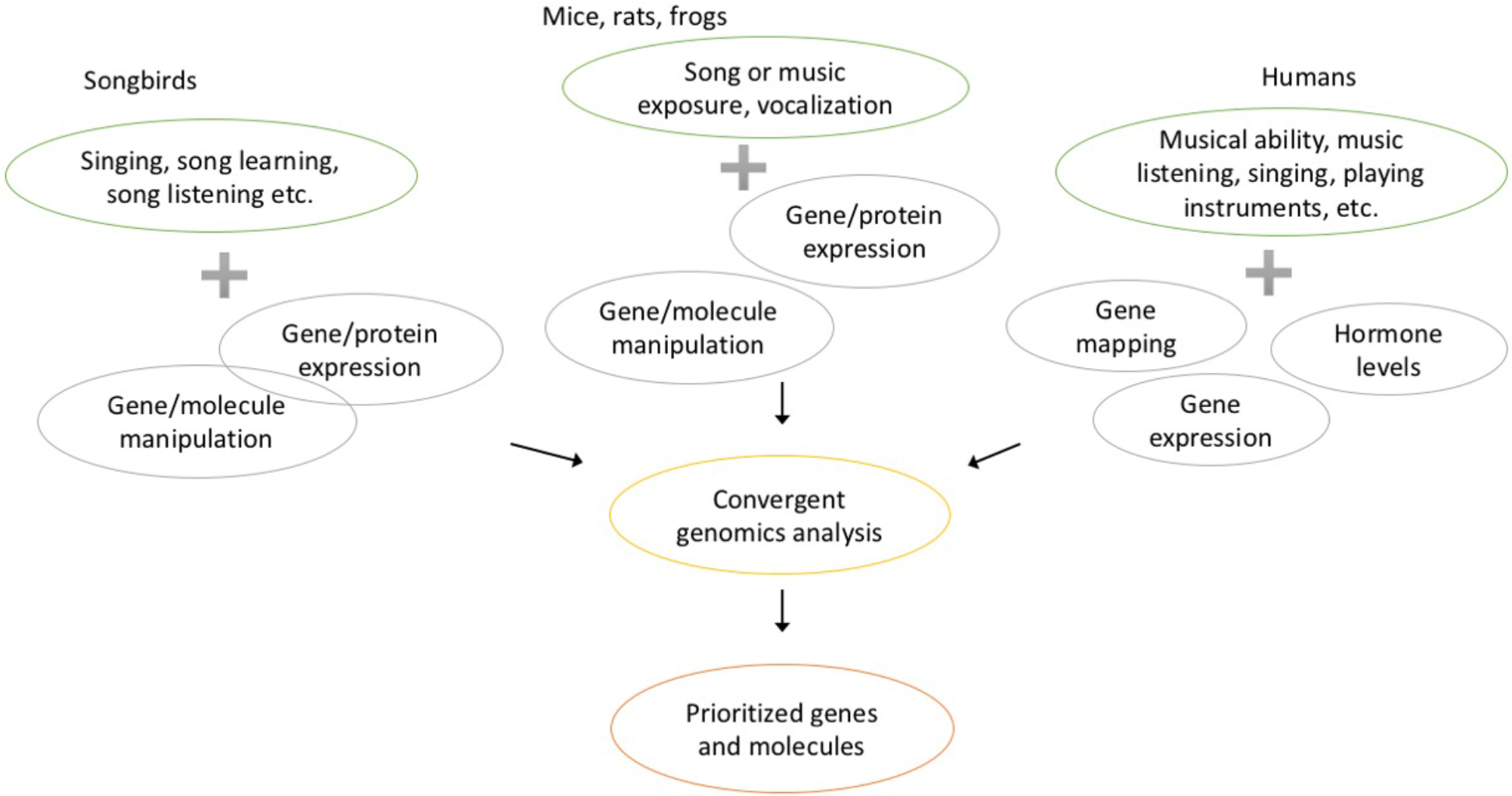
Studies included for the CE analysis included a variety of methods and animals. The music-related traits in studies using different animals varies as well as the used molecular evidence levels. All molecular evidence was combined in the CE analysis.

**Figure 2.**
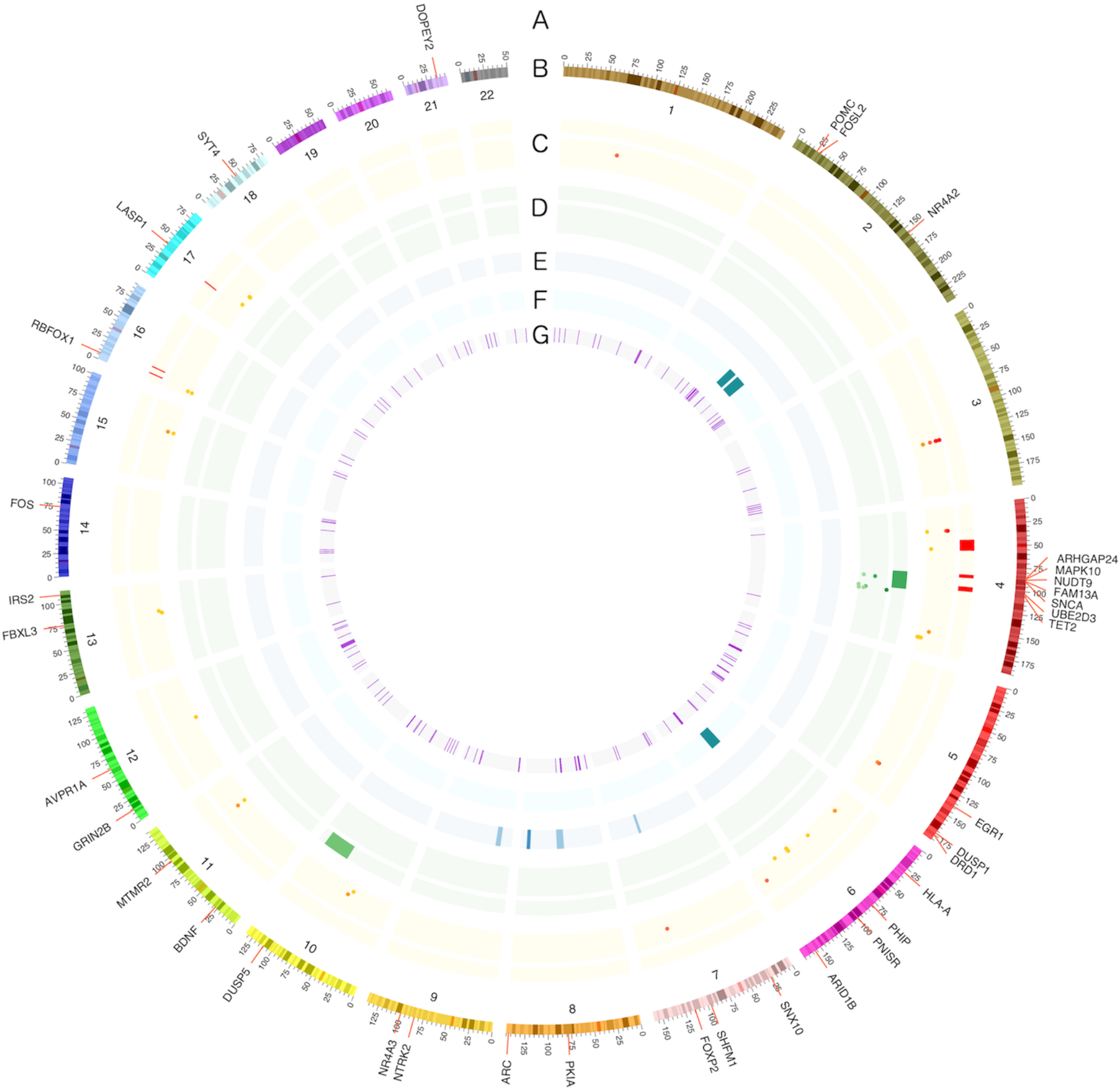
Human gene mapping results of musical abilities and the top 40 genes from the CE analysis. The top genes (A) are shown on the genomic context (B: coordinates for every chromosome as Mb). Published gene mapping results are shown with heat map bars (linkage) or dots (association): included results by (C) Oikkonen, et al. ^3^, (D) Park, et al. ^7^, (E) Theusch, et al. ^8^ and (F) Gregersen, et al. ^55^. The innermost circle shows regions identified by selection signature methods (G) by Liu, et al. ^13^

**Table 1.** Summary statistics of the study database. Numbers in braces in 3^rd^ and 4^th^ columns correspond to the number of genome-wide studies. Some studies include evidence from multiple molecular levels.

| Species | Number of studies | DNA,<br>genome-wide | RNA,<br>genome-wide | Other<br>levels | Sub-phenotypes |
| --- | --- | --- | --- | --- | --- |
| Human | 33 | 4 +(5) | 1+(3) | 20 | > 10 |
| Songbirds (> 10) | 55 | 0 | 16+(8) | 32 | > 17 |
| Other animals (> 4) | 13 | 0 | 1+(2) | 10 | > 5 |

### Top candidate genes

The CE method ranked *EGR1*, cortisol, *FOS*, *FOXP2*, *ARC*, dopamine and *BDNF* as the top candidate genes and molecules related to music (**Table 2, Supplementary Data**). Among the top hits, some genes like *EGR1*, *FOXP2, BDNF* and *ARC* have not yet been found in human studies, while some other genes like *PHIP, MAPK10, SNCA,* and *ARHGAP24,* received top ranks because of major evidence from human studies. The top candidate genes were evenly distributed across genomic locations, except for an enrichment of seven genes on chromosome 4q21-q24 (*MAPK10, SNCA, ARHGAP24, TET2, UBE2D3, FAM13A* and *NUDT9*), the region indicated in several gene mapping studies.

**Table 1.**
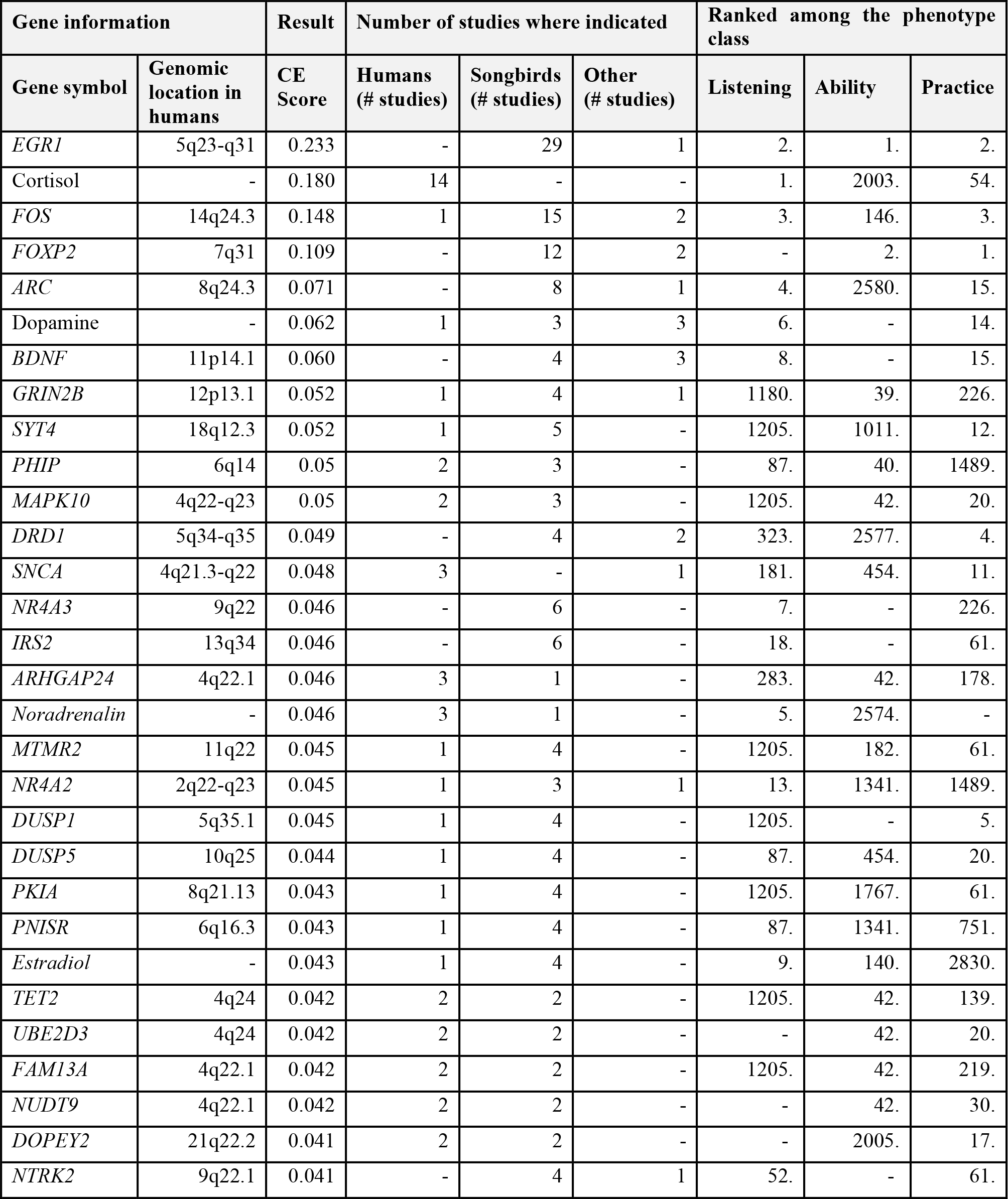
**Top 30 candidate genes related to music identified by CEReferences**

| Gene information |  | Result | Number of studies where indicated |  |  | Ranked among the phenotype class |  |  |
| --- | --- | --- | --- | --- | --- | --- | --- | --- |
| Gene symbol | Genomic location in humans | CE Score | Humans (# studies) | Songbirds (# studies) | Other (# studies) | Listening | Ability | Practice |
| *EGR1* | 5q23-q31 | 0.233 | - | 29 | 1 | 2. | 1. | 2. |
| Cortisol | - | 0.180 | 14 | - | - | 1. | 2003. | 54. |
| *FOS* | 14q24.3 | 0.148 | 1 | 15 | 2 | 3. | 146. | 3. |
| *FOXP2* | 7q31 | 0.109 | - | 12 | 2 | - | 2. | 1. |
| *ARC* | 8q24.3 | 0.071 | - | 8 | 1 | 4. | 2580. | 15. |
| Dopamine | - | 0.062 | 1 | 3 | 3 | 6. | - | 14. |
| *BDNF* | 11p14.1 | 0.060 | - | 4 | 3 | 8. | - | 15. |
| *GRIN2B* | 12p13.1 | 0.052 | 1 | 4 | 1 | 1180. | 39. | 226. |
| *SYT4* | 18q12.3 | 0.052 | 1 | 5 | - | 1205. | 1011. | 12. |
| *PHIP* | 6q14 | 0.05 | 2 | 3 | - | 87. | 40. | 1489. |
| *MAPK10* | 4q22-q23 | 0.05 | 2 | 3 | - | 1205. | 42. | 20. |
| *DRD1* | 5q34-q35 | 0.049 | - | 4 | 2 | 323. | 2577. | 4. |
| *SNCA* | 4q21.3-q22 | 0.048 | 3 | - | 1 | 181. | 454. | 11. |
| *NR4A3* | 9q22 | 0.046 | - | 6 | - | 7. | - | 226. |
| *IRS2* | 13q34 | 0.046 | - | 6 | - | 18. | - | 61. |
| *ARHGAP24* | 4q22.1 | 0.046 | 3 | 1 | - | 283. | 42. | 178. |
| *Noradrenalin* | - | 0.046 | 3 | 1 | - | 5. | 2574. | - |
| *MTMR2* | 11q22 | 0.045 | 1 | 4 | - | 1205. | 182. | 61. |
| *NR4A2* | 2q22-q23 | 0.045 | 1 | 3 | 1 | 13. | 1341. | 1489. |
| *DUSP1* | 5q35.1 | 0.045 | 1 | 4 | - | 1205. | - | 5. |
| *DUSP5* | 10q25 | 0.044 | 1 | 4 | - | 87. | 454. | 20. |
| *PKIA* | 8q21.13 | 0.043 | 1 | 4 | - | 1205. | 1767. | 61. |
| *PNISR* | 6q16.3 | 0.043 | 1 | 4 | - | 87. | 1341. | 751. |
| *Estradiol* | - | 0.043 | 1 | 4 | - | 9. | 140. | 2830. |
| *TET2* | 4q24 | 0.042 | 2 | 2 | - | 1205. | 42. | 139. |
| *UBE2D3* | 4q24 | 0.042 | 2 | 2 | - | - | 42. | 20. |
| *FAM13A* | 4q22.1 | 0.042 | 2 | 2 | - | 1205. | 42. | 219. |
| *NUDT9* | 4q22.1 | 0.042 | 2 | 2 | - | - | 42. | 30. |
| *DOPEY2* | 21q22.2 | 0.041 | 2 | 2 | - | - | 2005. | 17. |
| *NTRK2* | 9q22.1 | 0.041 | - | 4 | 1 | 52. | - | 61. |

To gather information whether the top genes relate only to a subset of the traits related to music, we analysed the studies in three phenotype categories: music listening, musical ability and music practice. The *EGR1* gene was among the highest ranking genes in all three categories, whereas a similar pattern was seen with none other molecules. Majority of the top genes were related to both music practice and listening (**Table 2, Supplementary Fig. S1**), with some exceptions: *PHIP,* noradrenalin and *NR4A2* were ranked among the top molecules in the whole sample as well as within music listening studies, but not within music practice (e.g. singing) related studies. Vice versa, genes like *DUSP1, PKIA* and *DOPEY2* were highly ranked in the whole sample as well as in music practice studies, but not in music listening or musical ability studies. Only a few top genes were most evident in the musical ability-related studies. These included the *GRIN2B* and *ARHGAP24* genes. Notably, studies related to each of the subphenotypes have also differences regarding the type of the biological samples and utilized animals: for example, there are more DNA-level and human studies relating to musical ability than to music listening or music practice (**Supplementary Data**).

### Functional annotation

We studied the functional annotations of the top ranked genes through functional class enrichment, pathway and interaction network analysis. Functional enrichment analysis of the top 40 ranked genes revealed enrichment of genes related to cognition, learning, memory, excitation of neurons, quantity of catecholamine, apoptosis of neurons, and long-term potentiation (p-values < 3^*^10^−10^), in the order of significance (p-value). ~ 40% of the top candidate genes were related to cognition (**Supplementary Table S5, Supplementary Fig. S2**). Pathway analysis showed enrichment of genes related to CDK5 signaling pathway (p-value 2.9^*^10^−8^, 6 molecules, **Supplementary Fig. S3**) that affects neurite outgrowth and neuronal migration^26^.

Further, network analysis revealed enrichment of interactions forming four non-overlapping networks each including 5-11 of the top-ranked genes. The most significant network was related to behaviour (Supplementary **Fig. S4**), the biological impact of the other three networks was less clear (**Supplementary Fig. S5-S7**). These networks may reveal subsets of the top molecules that work together. Because four non-overlapping networks were detected, they can indicate several separate pathways affecting music-related traits. For example, most of the genes within the second largest network (**Supplementary Fig. S6**) were indicated in studies considering music practice. Interestingly, also the genes related to CDK5 signaling pathway were enriched to the best, behaviour-related interaction network.

Upstream regulator analysis suggested cocaine, noradrenalin and calcineurin as possible upstream regulators (p-value < 6*10^−12^) of many of the top ranked genes. This shows interconnections between the top ranked genes because noradrenalin itself was one of the top molecules. Noradrenalin is known to increase expression of *BDNF, DUSP1, EGR1, FOS, FOSL2, IRS2, NR4A3* and *POMC* and decrease expression of *GRIN2B*, identified in this study. It works as a hormone and neurotransmitter causing arousal, anxiety and attention. Emotional arousal is known to affect memory and synaptic plasticity and noradrenalin is a major transmitter of these effects^27^.

## Discussion

We were able to identify 101 molecular studies including music-related phenotypes. We prioritized the genes and molecules found in these studies to identify the most probable genes affecting musical ability and the effects of music. Overall, *EGR1* was the highest ranked gene. General cognition-related genes were enriched within the top ranked genes.

Musical abilities and several related traits have been studied in humans and multiple animal models using genetic methods ranging from single-gene to genome-wide studies. Over the past decade, numerous genomic regions, genes and biomarkers related to musical abilities, including for example music perception and performance, have been detected with varying levels of statistical significance. Here, for the first time, we obtained aggregate evidence for the molecular basis of musical traits that are shared between species.

Several of the top candidate genes like *EGR1, FOS, ARC, BDNF* and *DUSP1* are activity-dependent immediate early genes (IEGs) well known to be regulated by sensory and motor behaviours in the brain^28,29^. These genes have been found to have essential roles in vocal learning and sound production in several animal studies^28,29^. Especially the *EGR1* gene has repeatedly been shown to upregulate during song listening and singing in zebra finches and other songbirds (see for example Avey, et al. ^30,^Drnevich, et al. ^31,^Jarvis, et al. ^32,^Mello, et al. ^33^). Interestingly, *FOS* and *DUSP1* genes have an increased transcriptional activity in professional musicians after they played musical instruments^11^. Other candidate genes like *FOXP2* and *GR1N2B* have been shown to be critical for vocal communication in songbirds and speech in humans^34–37^. Moreover, *GR1N2B* is located in positive selection region of musical aptitude^13^. Some of the top candidate genes have major evidence from animal studies (especially songbirds), with little or no evidence from human studies. The reason could depend on several factors like tissue-specificity, species-specificity and phenotype definition. It is important to note that the majority of animal studies were carried out on brain tissue, while it is mostly inaccessible in humans. Although we are interested in the shared molecular mechanisms behind musical abilities between species, we acknowledge that extrapolating genes found from animal models to human phenotypes carry limitations.

Among the top 40 candidate genes, seven genes (*MAPK10, SNCA, ARHGAP24, TET2, UBE2D3, FAM13A* and *NUDT9*) are located on chromosome 4q21-q24 (**Figure 2**), the region showing strongest evidence for linkage and association with musical abilities^3,7,38^. All these seven genes show major evidence from human studies, while receiving only supporting evidence from animal studies (Table 2). The other genes like *PHIP, DOPEY2, AVPR1A* and *POMC* that have been detected in human studies have not shown very strong evidence in individual studies. For example, *PHIP* has been shown only in one linkage analysis and one association analysis including humans but it has also been indicated in songbird-studies for singing and song listening^31,39^. However, our aggregated evidence now prioritizes these as the top candidate genes for musical abilities and effects of music. This demonstrates the strength of the CE method to prioritize candidate genes from multiple independent studies, even when some studies may be limited by the statistical power. On the other hand, genes like *UGT8, GATA2* and *PCDH7*, which were the most statistically significant findings in previous human genome-wide linkage and association studies, were not ranked among the top candidate genes here. This is probably due to the inherent nature of the method to identify only the genes with multiple independent evidence. Similarly, previous evidence suggested a role for auditory pathway genes in musical abilities, but this was not supported by the top ranked genes. This may result from the lack of music-related expression data from the inner ear and most other hearing-related tissues in this study, as well as from the wide spectrum of phenotypes included, not all related to perceptional skills.

Musical abilities (irrespective of the species) include the abilities to recognize, memorize and produce sound, which requires higher cognitive functions like learning and memory. Thus, the enrichment of candidate genes related to these functions in our study was not surprising. Moreover, an abundance of neuroscientific studies has demonstrated enhanced cognitive performance, learning and memory after training music for longer time^40–42^. Further, the enrichment of genes related to neuronal excitation, neuronal apoptosis and long-term potentiation support the idea that these neuronal plasticity-related physiological mechanisms are essential in music^43^. Cognitive functions like learning and memory require CDK5 regulation in the brain^44^, thus the enrichment of CDK5 signaling pathway-related genes in our study may explain the molecular basis of these cognitive processes involved in musical abilities. In songbird studies, MEK, which is a part of CDK5 signaling pathway (**Supplementary Fig. S3**), is necessary for song learning^45^. However, the CDK5 signaling pathway includes only a few genes identified in human studies of musical traits. The use of animal models for the study of musical traits enables the detection of this kind of brain-specific pathways that cannot be directly shown with human expression studies.

Although it has been suggested that genes like *EGR1* may relate to complex social structures and social communication instead of plain auditory stimuli in songbirds^46,47^, it may relate partially to both, speech and music. In human studies, it has been shown that musically trained individuals perform better in speech discrimination^48^. Also, *FOXP2* related genetic mutation has been found in a family with language and rhythm impairment^49,50^. Thus, partially similar features and genes are needed in music and in speech.

In the CE analysis, we included all identified molecular animal and human studies related to music, including singing, music playing and listening, musical abilities and vocal learning related traits. The analysis ranked highest those genes, which show evidence from multiple studies. The chosen layering ensured sensitivity for molecules, whereas specificity for musical traits was not prioritized. However, the combination of all these studies with varying phenotypes may reveal genes that are shared between most music-related traits.

The genetic predisposition for musical abilities is partially shared with general cognition^2,51^, which was also evident by the enrichment of cognition-related genes among the top candidate genes in this study. As the cognitive capacities in humans have undergone rapid evolution, the musical abilities - related genes or pathways might have also evolved more in humans than in other animals. Thus, there can be human-specific pathways and genes affecting musical abilities not captured by analyses in this work. For example, human-specific pathways have been shown to be important in Alzheimer disease^52^. Due to methodological differences, the human expression studies are enriched in the targets of the molecular pathways in music listening or practising whereas animal model studies mostly focus on the primary molecular effects in the brain. Similarly, DNA-level evidence on musical abilities was only obtained from humans. Hence, this study focuses on shared genetic pathways of musical abilities and music stimuli between animal species, while the special characteristics related to specific phenotypes will remain unresolved.

An obvious direction for future studies is to carry out phylogenetic analyses of the genes prioritized here: is there evolutionarily long-term conservation of the genes, or rather, convergent evolution? Also, it would be of interest to study the more specific role of the genes pinpointed: are some of them more related to the development of the brain, and some to the function in presence, as in reacting to vocal stimuli? The study was performed to elucidate key genes and novel pathways for music-related traits. The identified networks and pathways can guide future studies on genetic predisposition for music. Moreover, the gathered molecular information can be used to prioritize results in future studies considering music.

## Methods

### Study material

We collected studies related to music, which reported either genes or biomarkers. These included for example genetic linkage and association studies, expression and knockdown studies. The phenotypes in human studies included musical ability, absolute pitch, music listening, singing and playing instruments. In animal model studies the phenotypes were related for example to music and song exposure, vocal learning, singing and vocalization. Common denominators in the variable phenotypes were music perception and practise. The animal studies were searches to look for phenotypes that model these abilities.

The articles were collected with extensive searches through Google Scholar (https://scholar.google.fi), PubMed (http://www.ncbi.nlm.nih.gov/pubmed/) and Web of Science (http://apps.webofknowledge.com). We also searched articles from the references of the included studies and related reviews. Based on the titles, we chose a list of 326 studies that were further examined for relevance. Studies were excluded if the phenotype was not relevant to music, if there were no significant gene or biomarker-related results reported, or if there was a similar work from the same group already included (e.g. replicative studies from the same group with similar experimental settings). Results were extracted from a total of 101 short-listed articles using available data or by contacting the authors (**Supplementary Tables S1-S3**).

### Data extraction

All the extracted results were linked to corresponding human homologous genes using primarily biomaRt^53^ from Bioconductor. HGNC (HUGO Gene Nomenclature Committee) gene symbols were used when available^54^. Human association and linkage results were mapped into hg38 reference genome through markers (Ensembl, GRCh38.p5) and the genes within the regions were extracted. The linkage regions were identified from available data using the reported linkage regions when available. If only cM information was found, the regions around the reported peaks (usually nearest marker reported) were translated into Mb using 1cM = 1Mb formula. The association regions were identified as ±500kb around the associated markers. Some associated markers did drop out from the study as no genes were identified within the boundaries. To identify the associated markers, we used the thresholds reported in the studies. When there were significant and suggestive results reported in gene mapping studies, we included both of them because the convergent evidence method benefits from the integration of a larger number of results. From our gene mapping study, we included all association results above probability score 0.2 and linkage above 0.3^3^.

The human homologs for the bird and other species genes were gathered from the Ensembl BioMart data mining tool (http://www.ensembl.org/biomart/martview), and in the instances where they could not be accessed with BioMart, data was gathered using the Ensembl (http://www.ensembl.org), UniProt (http://www.uniprot.org), miRBase (http://mirbase.org, version 21), EggNOG (http://eggnogdb.embl.de), OrthoDB (http://cegg.unige.ch), BLAST (http://blast.ncbi.nlm.nih.gov/Blast.cgi) and Bird Base (http://birdbase.arizona.edu) databases. If there were multiple homologs per gene, all of them were included. Most of the genes that were found from the database were matched successfully to human homologs. However, there were a maximum of 20% of the resulting probes in some microarray studies where no current gene information was found.

The reported proteins were translated into genes encoding them. Hormones and other biomarkers that are synthesized from other substances, like estradiol, cortisol and dopamine, were included as such. Thus, the final data included genes and biomarkers.

### Convergent evidence

We used the convergent evidence method implemented in GenRank Bioconductor package (http://bioconductor.org/packages/GenRank) to prioritize candidate genes of musical traits. This method ranks genes by integrating gene-level data from multiple evidence layers. The rank of a gene depends upon the self-importance of each evidence layer it has been detected in and the number of evidence layers it has been detected in (**Supplementary Fig. S8**). The studies were input as separate layers and multiple evidence layers within gene mapping studies also as separate layers. Additionally, reanalysed transcriptome datasets in a study by Drnevich, et al. ^31^ were each treated as separate layers. This resulted in 105 evidence layers from the 101 articles in the analysis.

### Custom scores

We assigned differential scores to each evidence layer based upon the self-importance of each evidence layer. The self-importance of each evidence layer was determined based on three arguments: sample size, phenotype and homology conversion. These three arguments were each scored from 0.8 to 1 and multiplied to form a final score for each evidence layer. The idea behind this differential scoring strategy is to penalize those evidence layers that are limited by sample size, definition of phenotype and possible errors in homology conversion. The studies that precisely used musical traits as the phenotypes were given score 1 for the phenotype and scoring for all other related phenotypes was reduced to 0.8. Human-related studies were emphasized with full scoring for homology conversion while studies of other species were given 0.8 as the homology conversion in animal models may contain errors. Also, some genes may not be found or they may have different function. The sample sizes were given continuous scoring from 0.8 to 1 within two groups of studies: gene mapping studies and other studies (including for example protein level, hormone, gene expression and knockout studies). Within both study groups, the sample sizes were linearly scored from the smallest (getting 0.8) to the largest (getting 1.0). Additionally, the linkage layers were given reduced scoring (score multiplied by 0.9) if there were both linkage and association included from the same study. Therefore, final scores ranged from 0.512 to 1.

Studies were further divided into three phenotype classes related to music listening, musical ability and music practice. These classes were also separately analysed to find possible differences between the subphenotypes.

Enrichment analyses of biological functions were performed for the top ranked genes through the use of QIAGEN’s Ingenuity Pathway Analysis (IPA, QIAGEN Redwood City, http://www.ingenuity.com). We performed enrichment analyses of pathways, functional classes and upregulators to search for enriched biological functions among the top genes. Additionally, interaction network analysis was performed to study interconnectivity between the top genes. We chose top 40 genes for these analyses.

## Acknowledgements

The work is supported by the Academy of Finland (#13771), the Biomedicum Helsinki Foundation, the Finnish Brain Foundation, the Finnish Concordia Fund and the University of Helsinki.

The authors declare no conflict of interest.

## Author Contributions

J.O. participated in the design of the study, collected the material, carried out the statistical and bioinformatics data analyses and interpretation and drafted the manuscript.

P.O. participated in the design of the study and drafted the manuscript.

I.J. participated in the design of the study, collection of the material and drafted the manuscript.

C.K. conceived the idea of the study, participated in the collection of the material, performed part of the bioinformatics analyses and drafted the manuscript. All authors reviewed the manuscript.

